# Callosal fiber length scales with brain size according to functional lateralization, evolution, and development

**DOI:** 10.1101/2021.04.01.437788

**Authors:** Liyuan Yang, Chenxi Zhao, Yirong Xiong, Suyu Zhong, Di Wu, Shaoling Peng, Michel Thiebaut de Schotten, Gaolang Gong

**Affiliations:** State Key Laboratory of Cognitive Neuroscience and Learning & IDG/McGovern Institute for Brain Research, Beijing Normal University, Beijing, China; Brain Connectivity and Behaviour Laboratory, Sorbonne Universities, Paris, France; Groupe d’Imagerie Neurofonctionnelle, Institut des Maladies Neurodégénératives-UMR 5293, CNRS, CEA, University of Bordeaux, Bordeaux, France; Beijing Key Laboratory of Brain Imaging and Connectomics, Beijing Normal University, Beijing, China; Chinese Institute for Brain Research, Beijing, China

**Author notes:** **Correspondence:** Dr. Gaolang Gong.

## Abstract

Brain size significantly impacts the organization of white matter fibers. Fiber length scaling – the degree to which fiber length varies according to brain size – was overlooked. We investigated how fiber lengths within the corpus callosum, the most prominent white matter tract, vary according to brain size. The results showed substantial variation in length scaling among callosal fibers, replicated in two large healthy cohorts (∼2000 individuals). The underscaled callosal fibers mainly connected the precentral gyrus and parietal cortices, whereas the overscaled callosal fibers mainly connected the prefrontal cortices. The variation in such length scaling was biologically meaningful: larger scaling corresponded to larger neurite density index but smaller fractional anisotropy values; cortical regions connected by the callosal fibers with larger scaling were more lateralized functionally as well as phylogenetically and ontogenetically more recent than their counterparts. These findings highlight an interaction between interhemispheric communication and organizational and adaptive principles underlying brain development and evolution.

**Significance Statement:** Brain size varies across evolution, development, and individuals. Relative to small brains, the neural fiber length in large brains is inevitably increased, but the degree of such increase may differ between fiber tracts. Such a difference, if it exists, is valuable for understanding adaptive neural principles in large versus small brains during evolution and development. The present study showed a substantial difference in the length increase between the callosal fibers that connect the two hemispheres, replicated in two large healthy cohorts. Altogether, our study demonstrates that reorganization of interhemispheric fibers length according to brain size is intrinsically related to fiber composition, functional lateralization, cortical myelin content, evolutionary and developmental expansion.

## Introduction

Total brain size varies dramatically across evolution, development, and individuals. Large brains’ structures are not simple linearly scaled versions of small brains, e.g., cortical expansion of large brains is disproportionately larger for higher cognitive function regions (Hill et al., 2010; Reardon et al., 2018). The scaling patterns of specific brain structures provide essential clues for understanding organizational and adaptive principles underlying brain development and evolution (Zhang and Sejnowski, 2000; Herculano-Houzel et al., 2010; Van Essen, 2018).

An increase in brain size also has a significant impact on brain circuits (Kaas, 2000). In larger brains, fiber length is inevitably increased, accompanied by a longer delay in conduction and slower information transfer. It is well known that brain fibers can increase their myelination and diameter to increase conduction velocity and compensate for this delay (Waxman et al., 1995; Wang et al., 2008). These adaptations, however, cannot apply systematically to all fiber tracts due to the limit in brain space and energy consumption (Ringo et al., 1994), leading to alternative biological strategies that might impact fiber length differently. While the fiber length adjustment to brain size has been assumed uniform so far, heterogeneity in fiber length scaling may reveal adaptive principles underlying brain development and evolution.

In the present study, we assessed whether length scaling variations exist according to brain size within the corpus callosum (CC), the principal white matter (WM) bundle supporting the communication between the two brain hemispheres. We further investigated the relationship between CC length scaling with fiber composition, functional lateralization, cortical myelin content, and cortical evolutionary and developmental expansion.

## Materials and Methods

### Discovery dataset

The discovery dataset included all possible Human Connectome Project (HCP) participants for whom both diffusion and T1 images were available (Van Essen et al., 2012). It comprised 928 healthy young adults (female/male: 503/425; age range: 22-37 years). Retest diffusion and T1 imaging data were available on 35 of these subjects (test-retest interval: 1 month - 11 months). Informed consent was obtained from all subjects, and the protocol was approved by the Institutional Review Board of Washington University. Magnetic resonance imaging (MRI) scanning was performed using a customized Siemens Connectome Skyra 3T scanner. Diffusion-weighted (DW) images were acquired using a spin-echo echo-planar imaging (EPI) sequence with the following parameters: repetition time (TR) = 5520 ms, echo time (TE) = 89.5 ms, flip angle = 78°, field of view (FOV) = 210 × 180 mm^2^, matrix = 168 × 144, slices = 111, and resolution = 1.25 mm × 1.25 mm × 1.25 mm. Diffusion weightings of b = 0, 1000, 2000, and 3000 s/mm^2^ were applied in 18, 90, 90 and 90 directions, respectively. High-resolution 3D T1-weighted (T1W) images were acquired using a magnetization prepared rapid gradient echo (MPRAGE) sequence and the following parameters: TR = 2400 ms, TE = 2.14 ms, TI = 1000 ms, flip angle = 8°, FOV = 224 × 224 mm^2^, and resolution = 0.7 mm × 0.7 mm × 0.7 mm. DW and T1W images were preprocessed using the HCP minimal preprocessing pipeline (Glasser et al., 2013).

### Replication dataset

A subset of the UK Biobank (UKB) dataset was selected as our replication dataset (Miller et al., 2016). To maximally match the age range and sample size with the discovery dataset, all UKB participants under the age of 43 years for whom both diffusion and T1 images were available were included in the replication dataset, comprising 981 healthy adults (female/male: 562/419; age range: 40-43 years). Informed consent was obtained from all UKB subjects, and the protocol was approved by the North West Multicenter Research Ethics Committee. MRI data from the UKB sample were acquired using a Siemens Skyra 3T scanner. DW images were acquired using a spin-echo EPI sequence with the following parameters: TR = 3600 ms, TE = 92 ms, flip angle = 78°, FOV = 208 × 208 mm^2^, 72 slices, and resolution = 2 mm × 2 mm × 2 mm. Diffusion weightings of b = 0, 1000, 2000 s/mm^2^ were applied in 5, 50, and 50 directions. High-resolution 3D T1W images were acquired using an MPRAGE sequence and the following parameters: TR = 2000 ms, TE = 2.01 ms, TI = 880 ms, flip angle = 8°, FOV = 256 × 256 mm^2^, and resolution = 1 mm × 1 mm × 1 mm. DW and T1W images were preprocessed using the UKB brain imaging processing pipeline (Alfaro-Almagro et al., 2018).

### Total brain size

The brain size was defined as the total brain volume (TBV). Specifically, the T1W image of each individual was segmented into gray matter (GM), WM, and cerebrospinal fluid (CSF) using the FAST tool of the FMRIB Software Library (FSL) (Smith et al., 2004). The TBV was then calculated as the sum of WM and GM volume.

### Individual midsagittal CC (mCC) and its alignment to the template

The HCP minimal preprocessing pipeline aligned the anterior commissure (AC), AC– posterior commissure (PC) line, and interhemispheric plane of the T1W images to the MNI template using a rigid transform of 6 degrees of freedom. This transform maintains the original size and shape of the brain. We applied the same aligning procedure to the T1W images of UKB participants. On the resultant T1W image of each individual, the midsagittal plane’s sagittal slice was selected, and the midsagittal CC (mCC) boundary was then manually outlined by a trained rater (D.W). The mCC boundary of 50 randomly selected participants was outlined two weeks later by the rater to assess the manual outlining reliability. The Dice coefficient of the mCC masks ranged from 0.95 to 0.99 (mean: 0.98, STD: 0.009), and the intraclass correlation coefficient (ICC) of the mCC area reached 0.99.

To ensure voxel-wise comparability across individuals, the individual mCC was nonlinearly aligned to a template mCC in MNI space, which was manually outlined on the MNI152 template image. Specifically, the 2D sagittal image of the template mCC was set as the target image (or the fixed image), and the 2D sagittal image of the individual mCC was set as the source image (or the moving image). The two 2D images were first smoothed with a 2D Gaussian smoothing kernel (sigma = 1) by using the ‘imgaussfilt’ function in MATLAB. First, we linearly aligned the source image to the target image (i.e., performed a 2D affine transform of 8 degrees of freedom, including translation, rotation, scaling, and shearing) using the ‘imregtform’ function in MATLAB. Next, we performed nonlinear registration from the linearly aligned source image to the target image using the demons algorithm that was implemented by the ‘imregdemons’ function in MATLAB (Thirion, 1998; Vercauteren et al., 2009). Here, three multiresolution image pyramid levels were used, and 100 iterations were estimated at each pyramid level. Gaussian smoothing (sigma = 1) was applied to regularize the accumulated displacement field at each iteration. To maximize the aligning accuracy, this whole nonlinear registration procedure was iterated 5 times. Finally, the linear and nonlinear displacement fields were successively applied to transform the individual mCC to the template space with linear interpolation. For each individual, the mCC alignment was carefully checked by visual inspection.

### Callosal fiber length

Diffusion magnetic resonance imaging (dMRI) tractography was used to extract callosal fiber streamlines passing through the mCC. First, fiber orientation distributions (FODs) of each voxel were estimated using multishell, multitissue constrained spherical deconvolution with a harmonic order of 8 and default parameters (Tournier et al., 2007; Jeurissen et al., 2014). Probabilistic fiber tracking was then performed using the 2nd-order integration over FODs (iFOD2) algorithm in Mrtrix3 (Tournier et al., 2019). Given our focus on callosal fibers, the mCC was set as an inclusion mask (i.e., saving only fibers traversing the mCC) in fiber tracking. Due to the ambiguous existence of callosal fibers connecting bilateral subcortical nuclei, all subcortical nuclei were set as exclusion masks (i.e., fibers traversing them were discarded). Here, subcortical nuclei, including the thalamus, hippocampus, amygdala, caudate, putamen, palladium, and accumbens, were extracted using the FIRST tool of the FSL (Smith et al., 2004). To improve the biological accuracy of the tractograms, the anatomically constrained tractography (ACT) framework was used, which incorporates prior anatomical information into the tractography (Smith et al., 2012). The anatomical information was obtained by segmenting T1W images into tissue partial volume maps (PVMs) for WM, GM, and CSF using the FSL tools. The resultant PVM images were then taken as anatomical priors into the ACT framework. For the HCP participants, the minimal preprocessed T1W images were well aligned with the DW images; therefore, the mCC and PVM images were already in the DW image space. For each UKB participant, we linearly aligned the fractional anisotropy (FA) image to the T1W images, and the inverted transform was then used to convert the mCC and PVM images to the DW image space. The detailed fiber-tracking parameters were as follows: step size = 0.625 mm (default), maximum curvature per step = 45°, FOD threshold = 0.05, and length range = 2.5-250 mm. For each subject, 10 million streamlines were generated by seeding from and ending into the ACT-generated GM-WM interface (GMWMI). For the UKB participants, the generated tractograms in the DW image space were transformed back into the T1W image space.

For each mCC voxel, the streamlines passing through it and linking the bilateral cerebral cortex were selected. Their average length was computed as the callosal length value for this voxel. One mCC map of fiber length was obtained for each subject, which was further transformed into the standard mCC template space described above. To evaluate the test-retest reproducibility of the voxel-wise measure for callosal fiber length, we reran the above analyses for the 35 HCP retest subjects with their retest diffusion and T1 imaging data (test-retest interval: 1 month - 11 months). For each mCC voxel, ICC was then calculated for the fiber length measure using the test-retest data.

### Length scaling coefficients with brain size

As was done previously (Reardon et al., 2018), we estimated the fiber length scaling coefficient with greater brain size using a multiple regression model: *log* (fiber length) = *β* * *log* (brain size) + *β_1_*\* age + *β_2_*\* sex + intercept, where the *β* represents the length scaling coefficient. In the model, age and sex were included as covariates. This regression model fitting was applied to each voxel of the mCC template, resulting in a mCC map for the callosal fiber length scaling coefficient (Fig. 1).

**Fig. 1.**
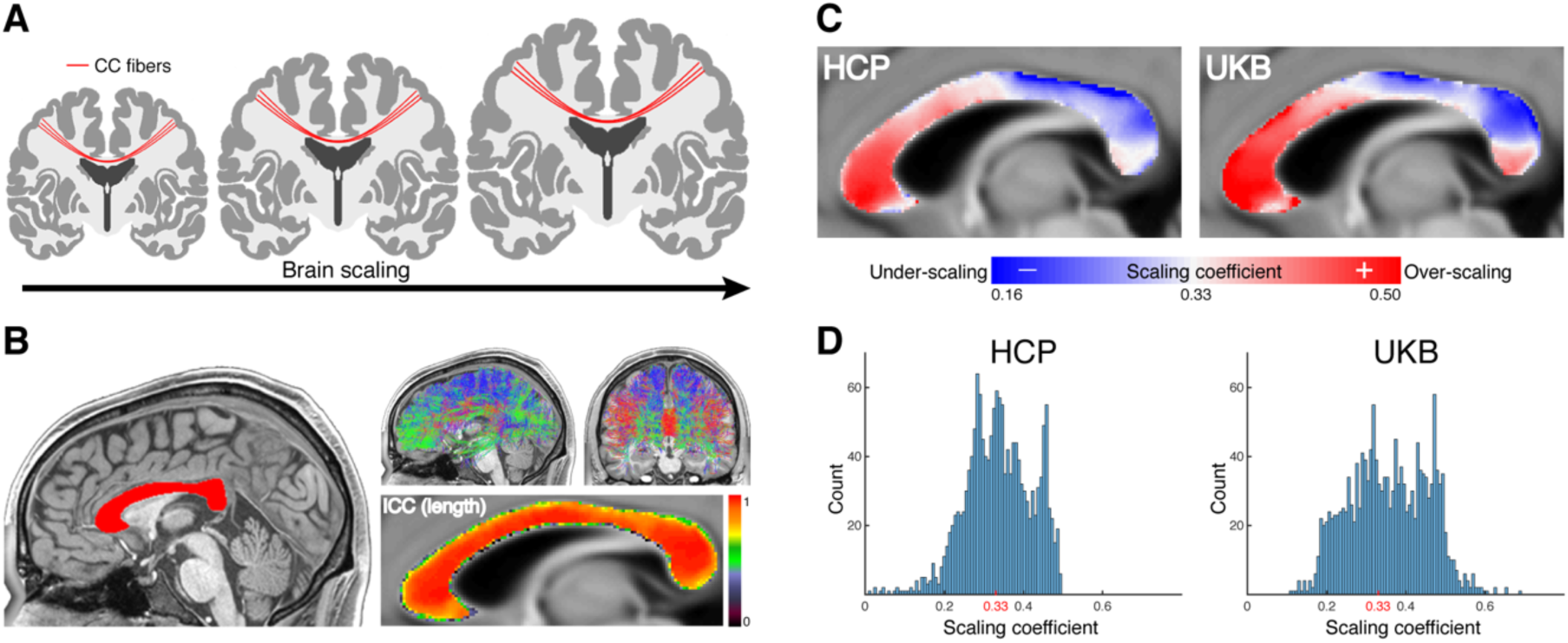
Callosal fiber variation in length scaling with brain size. (A) Schematic of callosal fiber length scaling with brain size. (B) The midsagittal CC (red) and callosal fibers from one example subject. The fiber length measure showed high intraclass correlation coefficient (ICC) values, suggesting excellent test-retest reproducibility. (C) Callosal fiber length scaling coefficient maps of the HCP and UKB datasets. (D) The histogram of scaling coefficients. The iso-scaling coefficient 0.33 is highlighted in red.

Theoretically, if WM fibers’ internal geometrical shape is invariant for brain size, the WM fiber length should scale as the 1/3 power of brain size. In this case, the proportional relationship between the two measures, one in 1-dimension and the other in 3-dimension, is preserved during scaling. This is referred to as the isometric scaling or iso-scaling. For empirical scaling coefficients, coefficients > 1/3 indicate that the WM fiber length scales more with greater brain size, i.e., over-scaling; coefficients < 1/3 indicate that WM fiber length scales less with brain size, i.e., under-scaling. Each voxel’s scaling coefficient on the mCC template was statistically compared to 1/3 to identify the significant clusters with over-scaling or under-scaling. To correct for multiple comparisons, the false discovery rate (FDR) procedure was performed at *q* value of 0.05 (Genovese et al., 2002).

### Validation analysis of callosal tractography

The validity of our callosal tractography is critical for measuring callosal fiber length and performing subsequent length scaling analyses. To validate our callosal tractography results, we evaluated a set of characteristics of tractography-based callosal connections as following.

First, we selected several cortical regions of interest (ROIs) that showed length under-scaling and over-scaling in scaling analyses, and investigated how tightly their callosal streamlines are clustered on the mCC and how the clusters on the mCC correspond with previous anatomical studies. Specifically, we extracted the cortical ROIs using the HCPMMP atlas (Glasser et al., 2016). The selected ROIs included 1) area 4 on the primary motor cortex, 2) area 2 on the somatosensory cortex, 3) area TE1p on the lateral temporal cortex, 4) area 7AL on the lateral parietal cortex, 5) area POS2 on the medial parietal cortex, 6) area 9-46d on the lateral prefrontal cortex, and 7) area 9m on the medial prefrontal cortex. For each ROI, a streamline count map was generated by projecting the ROI-connected callosal streamlines onto the mCC for each individual. The resultant maps were further binarized and averaged across individuals, resulting a group pattern map (Fig. 2A).

**Fig. 2.**
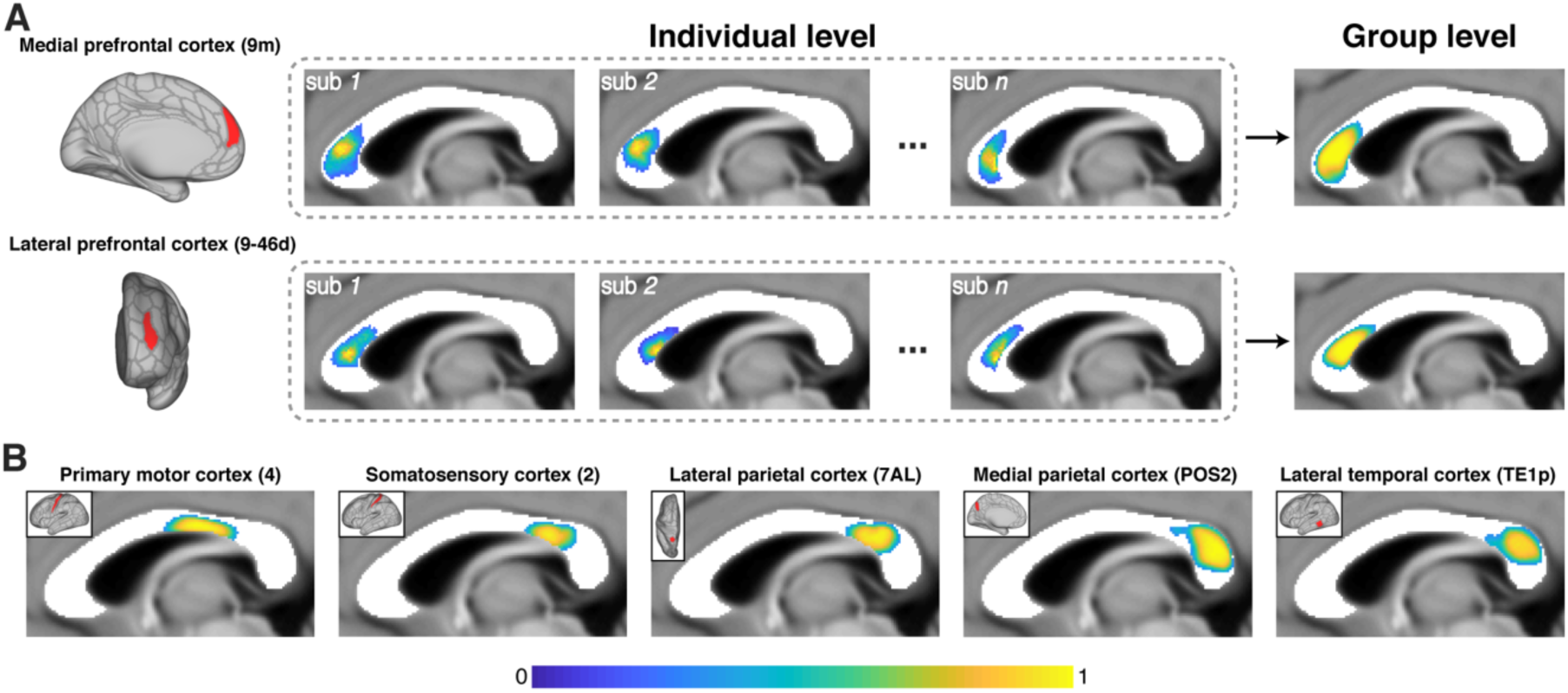
Callosal streamlines’ passing location on the mCC for selected cortical regions. (A) Schematic diagram of generating group mCC location for a cortical region. As illustrated, the passing location on the mCC can overlap to some degree for two different cortical regions (e.g., area 9-46d on the lateral prefrontal cortex and area 9m on the medial prefrontal cortex). (B) The resultant group mCC pattern maps are displayed for selected HCPMMP regions on the left hemisphere (red in the solid box): area 4 on the primary motor cortex, area 2 on the somatosensory cortex, area TE1p on the lateral temporal cortex, area 7AL on the lateral parietal cortex, and area POS2 on the medial parietal cortex. The HCPMMP regional abbreviation is included in the bracket.

For each cortical ROI above, we further evaluated the distribution of its callosal connections on the contralateral hemisphere. The connection strength between this ROI and HCPMMP regions on the contralateral hemisphere was measured by the fraction of streamlines (FSe), which is calculated as the fraction of streamlines linking two regions relative to the number of streamlines extrinsic to those regions (Donahue et al., 2016). The FSe is conceptually analogous to the connection strength in retrograde-tracer-based studies (Donahue et al., 2016), and has been increasingly used in recent studies (Zhao et al., 2019; Hayashi et al., 2021; Rosen and Halgren, 2021). Likewise, the group FSe map was obtained by averaging the FSe maps across individuals. For each ROI, the major connected contralateral regions, i.e., accounting for at least 80% of total FSe value from this region, were then extracted and displayed.

To assess how callosal streamline terminations in the contralateral hemisphere relate to cortical gyral and sulcal features, we measured the proportion of streamlines that were terminated on the cortical gyri.

For each mCC voxel, we also selected all passing callosal streamlines and calculated the number of streamlines that were terminated in each HCPMMP cortical regions in the left and right hemispheres, respectively. To quantify the similarity of the two hemispheric streamline number patterns, we computed two measures: Pearson correlation (*r*) and intraclass correlation coefficient (*ICC*), resulting an *r* map and an *ICC* map of mCC for each individual as well as group-averaged *r* and *ICC* maps of mCC.

Finally, while neither tractography-based structural connectivity (SC) nor fMRI-based functional connectivity (FC) is perfect, FC is more likely to reflect interhemispheric connectivity between homotopic regions in the two hemispheres. We therefore evaluated whether interhemispheric homotopic SC correlates with corresponding FC. To quantify the SC for each HCPMMP homotopic regional pair, we used the FSe as above. The group-averaged FSe was then generated and further log-transformed (Rosen and Halgren, 2021). Regarding the FC estimation, the two HCP rs-fMRI sessions were used: REST1(919 subjects; female/male: 495/424) and REST2 (894 subjects; female/male: 482/412). Specifically, we adopted the HCP minimally preprocessed rs-fMRI data, and the preprocessing procedures included magnetic gradient distortion correction, EPI distortion correction, nonbrain tissue removal, MNI standard space registration, and intensity normalization (Glasser et al., 2013). The resultant data were then denoised by using independent component analysis with the FIX tool (Salimi-Khorshidi et al., 2014). The procedures of linear detrending, global signal regression and temporal bandpass filtering (0.01-0.1 Hz) were further applied (Lowe and Russell, 1999; Cordes et al., 2001). For each HCPMMP region, the averaged rs-fMRI time series across all vertices was calculated as the regional rs-fMRI time series. For each HCPMMP homotopic regional pair, the FC strength was measured using the Pearson correlation coefficient (converted to Fisher’s Z-values) between the two regional time series. The group-averaged FC was then generated for each HCPMMP homotopic regional pair.

### WM microstructural measures on the mCC

To measure callosal fiber composition, two widely used dMRI-derived parameters, i.e., neurite density index (NDI) and fractional anisotropy (FA), were estimated for each mCC voxel (Basser et al., 1994; Zhang et al., 2012). The mCC maps of NDI and FA were further transformed to the standard mCC template space for each individual. The representative NDI and FA maps in healthy adults were then generated by averaging all HCP individual maps of the template space. Such representative maps were used to assess the relevance of length scaling to callosal fiber composition.

### Mapping cortical topography of each mCC voxel

For each HCP subject, the minimal preprocessing pipeline outputted FreeSurfer-generated pial and white surfaces resampled onto the standard 32k_fs_LR mesh in the native volume space. Each mCC voxel’s streamlines were assigned to their closest vertex on the white surface within a sphere with a 2-mm radius centered at its endpoint, therefore yielding a standard 32K surface map of streamline counts for this voxel. For each voxel on the mCC template, such a streamline count map on the standard 32K surface was comparable across individuals. The group streamline count surface map was then obtained by averaging the individual streamline count surface maps across all HCP individuals. We applied a threshold to the group streamline count surface map to extract a binary cortical topography map for each mCC voxel. Specifically, the mean + 0.5*standard deviation (STD) of the group streamline count value across the entire surface was used as the main threshold. To evaluate the influence of such a threshold on the results, we reran relevant analyses by applying the following two thresholds: 1) a less stringent threshold, i.e., the mean of the group streamline count value across the entire surface, and 2) a more stringent threshold, i.e., the mean + 1*STD of the group streamline count value across the entire surface.

### Cortical measures

To assess the relevance of callosal fibers’ length scaling to particular functional and structural aspects of their connected cortical regions, we derived several cortical measures; these measures are described below.

#### Functional lateralization index

We followed a recent study’s methods to estimate the cortical overall functional LI across cognitive domains (Karolis et al., 2019). First, we selected available cognitive terms (575 in total) in the current Neurosynth database (Yarkoni et al., 2011) (http://Neurosynth.org). We applied the Neurosynth tool for each cognitive term to generate a whole-brain meta-activation image in MNI space. After aligning this meta-activation image to a symmetric image template, we projected the values onto the standard 32k_fs_LR mesh in MNI space. For each surface vertex, the absolute between-hemisphere difference of the meta-activation values was calculated as the absolute degree of functional lateralization for this cognitive term. To yield an overall functional LI across all cognitive domains, we simply averaged the individual surface maps of functional lateralization across all 575 cognitive terms.

#### Myelin content index

The group-averaged cortical T1w/T2w ratio-based myelin content map, which was corrected for the transmit field bias, was estimated by Glasser and colleagues (Glasser et al., 2021). We obtained this map from https://balsa.wustl.edu/file/show/zpL2m, which is already on the standard 32k_fs_LR surface mesh.

#### Evolutionary expansion index

The cortical expansion map in humans relative to macaques was estimated by Hill and colleagues (Hill et al., 2010). We adopted the expansion index and resampled the values onto the standard 32k_fs_LR surface mesh.

#### Human developmental expansion index

The cortical expansion map in human adults relative to human infants was estimated by Hill and colleagues (Hill et al., 2010). Likewise, we adopted the expansion index and resampled the values onto the standard 32k_fs_LR surface mesh.

For each mCC voxel, we assigned the mean value of the aforementioned cortical measure within its connected cortical region, resulting in one mCC map for the cortical functional lateralization, cortical myelin content, evolutionary and developmental expansion.

### Statistical analysis

To determine whether the length scaling coefficient was related to callosal microstructural measures and cortical measures above, we calculated the Spearman correlations across all mCC voxels. We used the Spearman correlation because the length scaling coefficient showed a nonnormal distribution on the mCC. A permutation test (10,000 permutations) was adopted to estimate the Spearman correlation coefficient’s statistical significance (*r_s_*). To evaluate the robustness and reproducibility of the observed correlation across the analysis resolution of the mCC, we additionally parcellated the entire mCC into ten segments. The parcellation scheme followed an influential mCC study, which measured histological fiber density for each of the ten segments (Aboitiz et al., 1992). Likewise, we assigned the mean callosal microstructural measures or the mean cortical index value of the connected cortical region to each mCC segment. We then reran the Spearman correlation analysis across all ten mCC segments.

To examine an independent relationship between callosal fiber length scaling and the aforementioned cortical measures, we reran the Spearman correlations across the mCC voxels or segments described above while controlling for the NDI and FA.

## Results

### Validation of callosal tractography

As shown in Fig. 2, the streamlines from the primary motor cortex mainly traversed the posterior body; the streamlines from the somatosensory cortex mainly traversed the posterior body and anterior splenium; the streamlines from the lateral temporal cortex and medial parietal cortex were tightly distributed on the posterior splenium; the streamlines from the lateral parietal cortex were distributed on the anterior splenium. In contrast, the streamlines from the lateral and medial prefrontal cortex were mostly clustered on the genu. The observed spatial pattern on the mCC was strikingly similar for homotopic ROI pairs on the left (Fig. 2) and right (Fig. S1) hemispheres, indicating the robustness of the result. Importantly, these location patterns on the mCC are quite consistent with previous chemical tracing findings in monkey and histological observations in human (Barbas and Pandya, 1984; de Lacoste et al., 1985; Rockland and Pandya, 1986; Caminiti et al., 2009, 2013; Tomasi et al., 2012), therefore supporting the validity of our current CC tractography.

Fig. 3 illustrated the major callosal connections to regions on the contralateral hemisphere (i.e., accounting for at least 80% of total FSe value) for the selected cortical ROIs on the left hemisphere. As shown, the maximal FSe of the selected ROI was largely located in the homotopic contralateral region, and major connections to the contralateral hemisphere were concentrated around the homotopic and nearby regions. The only exception was the cortical ROI on the lateral temporal cortex, which likely relates to systematic biases of our current callosal tractography from the temporal lobe. The major callosal connections to the contralateral hemisphere for the selected cortical ROIs on the right hemisphere were illustrated in Fig. S2.

**Fig. 3.**
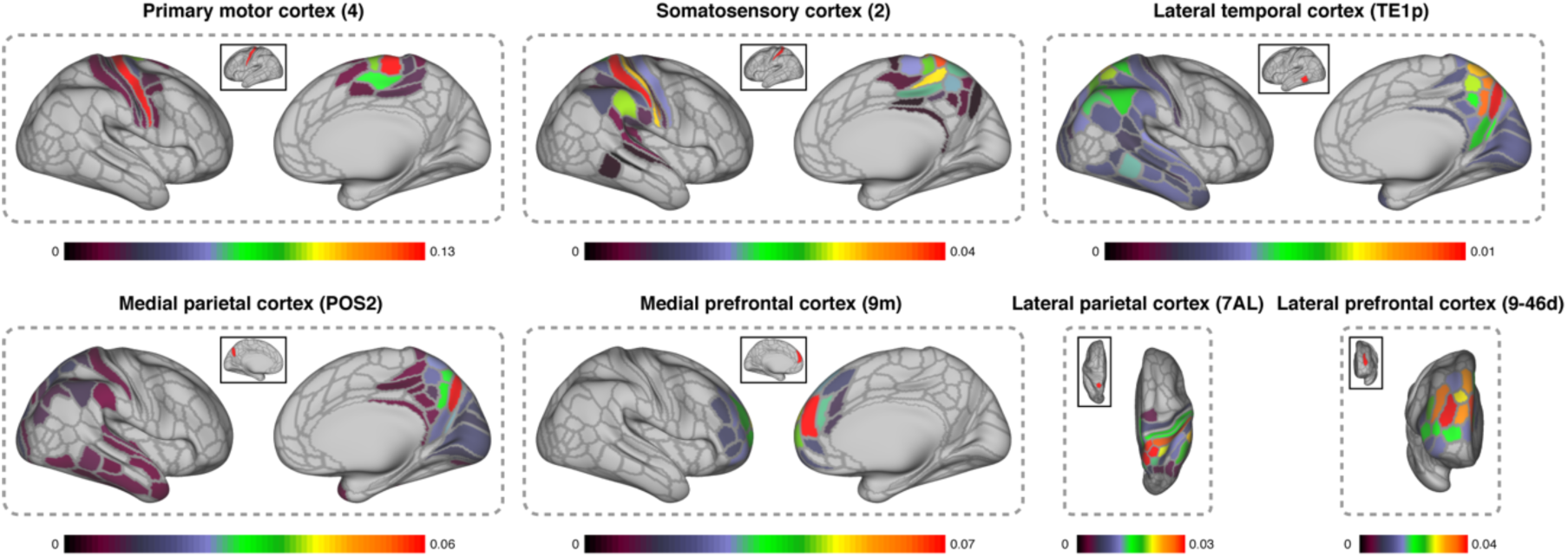
Major callosal connections to the contralateral hemisphere for selected cortical regions. For each cortical region, the top strongly connected regions on the contralateral hemisphere are displayed, which together account for at least 80% of total fraction of streamline (FSe) value from this region. The group FSe maps are displayed for selected HCPMMP regions on the left hemisphere (red in the solid box): area 4 on the primary motor cortex, area 2 on the somatosensory cortex, area TE1p on the lateral temporal cortex, area 7AL on the lateral parietal cortex, area POS2 on the medial parietal cortex, area 9-46d on the lateral prefrontal cortex, and area 9m on the medial prefrontal cortex. The HCPMMP regional abbreviation is included in the bracket. The color bar indicates the FSe value from the selected cortical region.

In line with previous anatomical observation (Xu et al., 2010; Nie et al., 2012; Budde and Annese, 2013; Chen et al., 2013), more gyrally terminated streamlines than sulcally terminated streamlines were observed in the vast majority of individuals (Fig. S3 and Fig. S4).

For all mCC voxels, their passing streamlines termination pattern on the hemisphere was quite similar between the two hemispheres (Fig. S5). This corresponds well with the largely mirrored nature of callosal connectional termination on the two hemispheres, therefore favoring the validity of our CC tractography.

However, we did not observe significant SC-FC correlation across the homotopic HCPMMP regional pairs (Fig. S6), which is consistent with a recent study (Rosen and Halgren, 2021). This negative result might reflect biases of our callosal tractography or biologically meaningful divergence between these two types of interhemispheric connectivity.

### Callosal fiber variation in length scaling with brain size

As shown in Fig. 1, the measure of voxel-wise callosal fiber length on the mCC showed excellent test-retest reproducibility (ICC: 0.83 ± 0.18). The callosal fiber length scaling coefficients showed substantial variation across mCC voxels. Specifically, the rostrum, genu, anterior body, and posterior splenium showed the largest scaling coefficients, whereas the smallest values were in the posterior body and isthmus parts of the CC. The mCC scaling coefficient maps estimated from both the HCP and UKB datasets were strikingly similar (Spearman correlation *r_s_* = 0.91, *p* < 10^-4^), indicating excellent reproducibility for such a scaling pattern across the mCC. We further re-evaluated the length scaling map by defining the voxel-wise fiber length as the median and mode value of all passing streamlines or by using the HCP unrelated samples to fit the log-log regression model. The resulting patterns remain highly similar (data not shown), indicating that the currently observed length scaling pattern on the mCC was also quite robust to the computing methods for voxel-wise fiber length and the selected samples for regression model fitting.

### Relevance to callosal fiber composition

To determine whether the length scaling relates to underlying fiber composition, we evaluated the Spearman correlation of the scaling coefficient with two popular dMRI-derived WM microstructural parameters on the mCC: NDI and FA. Across all mCC voxels, the length scaling coefficients were found to be positively correlated with NDI (HCP: *r_s_* = 0.43, *p* < 10^-4^; UKB: *r_s_* = 0.37, *p* < 10^-4^; Fig. 4A) and negatively correlated with FA (HCP: *r_s_* = -0.05, *p* = 0.03; UKB: *r_s_* = -0.12, *p* < 10^-4^; Fig. 4B). Moreover, we parcellated the mCC into Aboitiz’s ten segments, which measured histological fiber density for each of the ten segments (Aboitiz et al., 1992). Across the ten mCC segments, significant correlation was found between the histological fiber density of callosal fiber > 0.4µm and NDI (*r_s_* = 0.74, *p* = 0.008) but not with FA (*r_s_* = 0.12, *p* = 0.36). There was no significant correlation between the length scaling coefficient and NDI or FA (Fig. 4), possibly due to the limited statistical power.

**Fig. 4.**
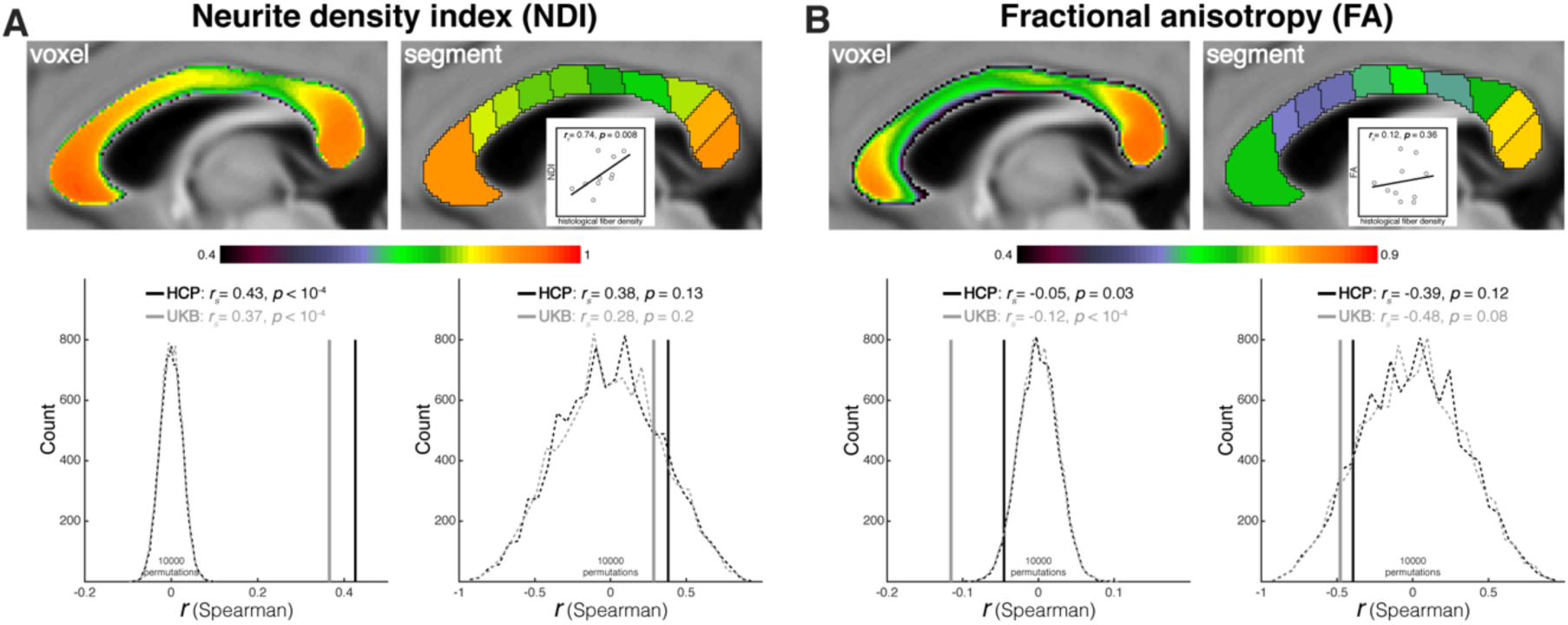
Relevance to callosal fiber composition. The relationships between callosal fiber length scaling and neurite density index (A), and fractional anisotropy (B). Top row is the group-averaged neurite density index (NDI) and fractional anisotropy (FA) at the voxel and segment level. The scatter plots of NDI and FA against the Aboitiz’s histological callosal fiber density are displayed at the segment level. Bottom row is the Spearman correlation between the microstructural measures and callosal fiber length scaling coefficient across midsagittal CC voxels and segments. Black: HCP, Gray: UKB. Solid lines represent empirical Spearman correlation coefficients; dashed lines represent the null distribution of Spearman correlation coefficients derived from 10,000 permutations.

### Relevance to cortical functional lateralization

The length scaling variation between callosal fibers putatively reflected cortical differences in evolutionary or developmental demands for rapid interhemispheric communication, which may have accounted for cortical differences in functional lateralization of the human brain. To assess this hypothesis, we estimated an overall functional lateralization index (LI) across cognitive domains (all 575 cognitive terms in the current Neurosynth database) on the cortical surface (Fig. 5A) using a Neurosynth-based meta-analysis (Yarkoni et al., 2011) (http://Neurosynth.org).

**Fig. 5.**
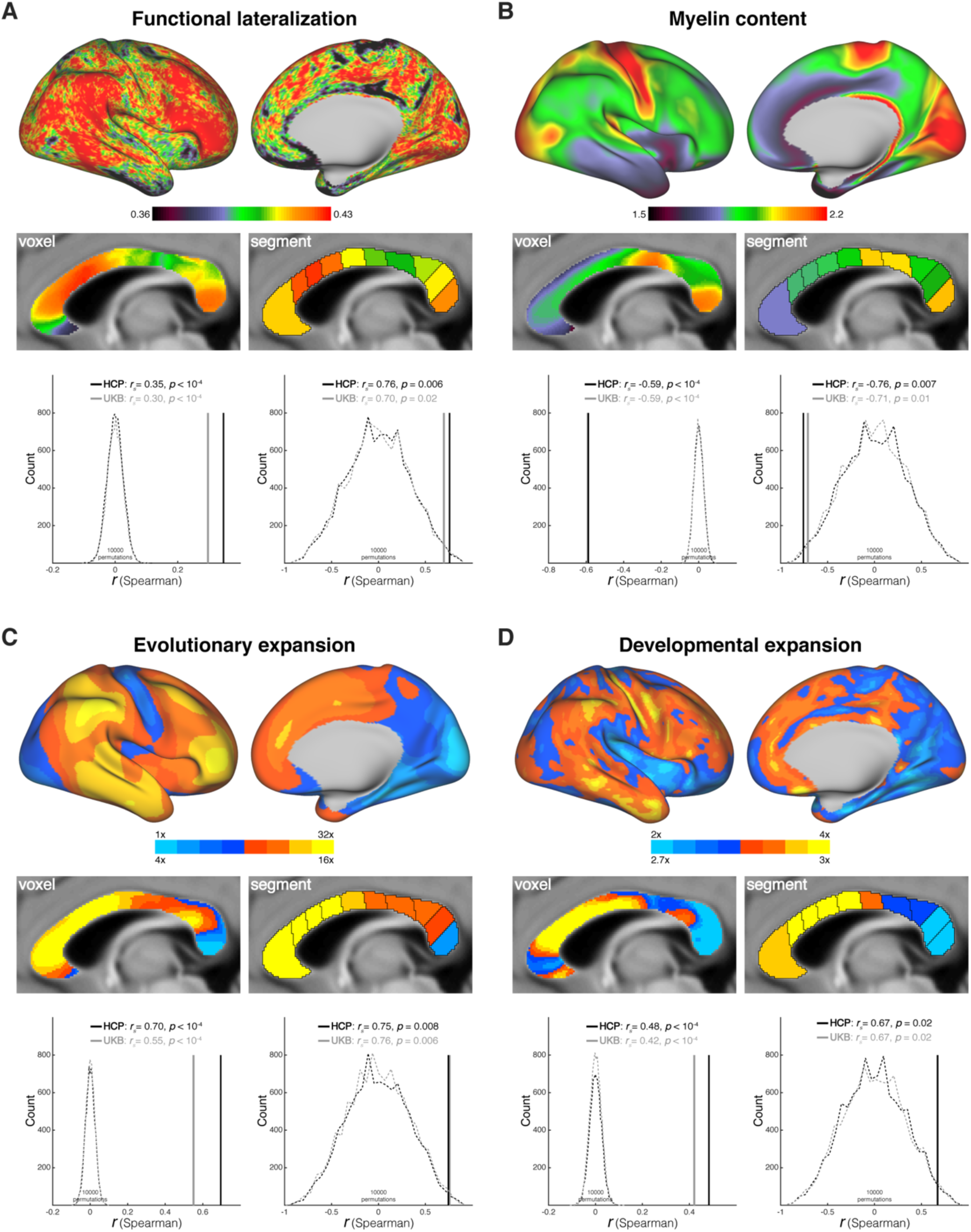
Relevance to cortical measures. The relationships between callosal fiber length scaling and cortical functional lateralization (A), myelin content (B), evolutionary cortical expansion of the primate brain (C), and developmental cortical expansion of the human brain (D). Top row is the averaged cortical functional lateralization map across cognitive terms, averaged myelin content map across HCP subjects, evolutionary cortical expansion map, and developmental cortical expansion map, respectively. Middle row is the cortical measures of the midsagittal CC at the voxel and segment level. Bottom row is the Spearman correlation between the cortical measures and callosal fiber length scaling coefficient across midsagittal CC voxels and segments. Black: HCP, Gray: UKB. Solid lines represent empirical Spearman correlation coefficients; dashed lines represent the null distribution of Spearman correlation coefficients derived from 10,000 permutations.

For each mCC voxel, we mapped its connected cortical region and calculated its regional mean functional LI. Across all mCC voxels, the length scaling coefficients were positively correlated with the functional LI (HCP: *r_s_* = 0.35, *p* < 10^-4^; UKB: *r_s_* = 0.30, *p* < 10^-4^; Fig. 5A). We also observed a positive correlation between length scaling and the functional LI across the ten Aboitiz’s segments (HCP: *r_s_* = 0.76, *p* = 0.006; UKB: *r_s_* = 0.70, *p* = 0.02), indicating the robustness of this correlation. Therefore, stronger cortical functional lateralization was accompanied by greater length scaling of the connected callosal fibers, thus suggesting less demand for rapid cortical interhemispheric communication. At both the voxel and segment levels, the partial Spearman correlations after controlling for the NDI and FA remained significant, indicating an independent relationship between callosal fiber length scaling and functional lateralization (Table 1).

**Table 1.** The full and partial Spearman correlation of the callosal fiber length scaling with cortical measures under different cortical mapping thresholds.

|  |  |  | Cortical mapping threshold |  |  |  |  |  |
| --- | --- | --- | --- | --- | --- | --- | --- | --- |
|  |  |  | Mean + 0.5*STD |  | Mean |  | Mean + 1*STD |  |
|  |  |  | HCP | UKB | HCP | UKB | HCP | UKB |
| Functional lateralization | Voxel | Full | 0.35 ($p<10^{-4}$)*** | 0.30 ($p<10^{-4}$)*** | 0.34 ($p<10^{-4}$)*** | 0.30 ($p<10^{-4}$)*** | 0.32 ($p<10^{-4}$)*** | 0.27 ($p<10^{-4}$)*** |
| | | Partial | 0.24 ($p<10^{-4}$)*** | 0.17 ($p<10^{-4}$)*** | 0.21 ($p<10^{-4}$)*** | 0.15 ($p<10^{-4}$)*** | 0.22 ($p<10^{-4}$)*** | 0.14 ($p<10^{-4}$)*** |
| | Segment | Full | 0.76 ($p=0.006$)** | 0.70 ($p=0.02$)* | 0.67 ($p=0.02$)* | 0.60 ($p=0.04$)* | 0.78 ($p=0.004$)** | 0.75 ($p=0.009$)** |
| | | Partial | 0.71 ($p=0.02$)* | 0.63 ($p=0.03$)* | 0.61 ($p=0.04$)* | 0.51 ($p=0.08$) | 0.74 ($p=0.01$)* | 0.68 ($p=0.02$)* |
| Myelin content | Voxel | Full | -0.59 ($p<10^{-4}$)*** | -0.59 ($p<10^{-4}$)*** | -0.64 ($p<10^{-4}$)*** | -0.64 ($p<10^{-4}$)*** | -0.53 ($p<10^{-4}$)*** | -0.55 ($p<10^{-4}$)*** |
| | | Partial | -0.23 ($p<10^{-4}$)*** | -0.21 ($p<10^{-4}$)*** | -0.29 ($p<10^{-4}$)*** | -0.26 ($p<10^{-4}$)*** | -0.14 ($p<10^{-4}$)*** | -0.16 ($p<10^{-4}$)*** |
| | Segment | Full | -0.76 ($p=0.007$)** | -0.71 ($p=0.01$)* | -0.82 ($p=0.002$)** | -0.71 ($p=0.01$)* | -0.79 ($p=0.003$)** | -0.71 ($p=0.01$)* |
| | | Partial | -0.70 ($p=0.02$)* | -0.61 ($p=0.05$)* | -0.78 ($p=0.008$)** | -0.61 ($p=0.04$)* | -0.75 ($p=0.01$)* | -0.62 ($p=0.04$)* |
| Evolutionary expansion | Voxel | Full | 0.70 ($p<10^{-4}$)*** | 0.55 ($p<10^{-4}$)*** | 0.64 ($p<10^{-4}$)*** | 0.52 ($p<10^{-4}$)*** | 0.69 ($p<10^{-4}$)*** | 0.55 ($p<10^{-4}$)*** |
| | | Partial | 0.64 ($p<10^{-4}$)*** | 0.37 ($p<10^{-4}$)*** | 0.53 ($p<10^{-4}$)*** | 0.32 ($p<10^{-4}$)*** | 0.64 ($p<10^{-4}$)*** | 0.37 ($p<10^{-4}$)*** |
| | Segment | Full | 0.75 ($p=0.008$)** | 0.76 ($p=0.006$)** | 0.65 ($p=0.02$)* | 0.60 ($p=0.04$)* | 0.75 ($p=0.007$)** | 0.70 ($p=0.01$)* |
| | | Partial | 0.70 ($p=0.02$)* | 0.67 ($p=0.02$)* | 0.56 ($p=0.06$) | 0.46 ($p=0.1$) | 0.73 ($p=0.01$)* | 0.58 ($p=0.05$) |
| Developmental expansion | Voxel | Full | 0.48 ($p<10^{-4}$)*** | 0.42 ($p<10^{-4}$)*** | 0.34 ($p<10^{-4}$)*** | 0.27 ($p<10^{-4}$)*** | 0.52 ($p<10^{-4}$)*** | 0.45 ($p<10^{-4}$)*** |
| | | Partial | 0.52 ($p<10^{-4}$)*** | 0.35 ($p<10^{-4}$)*** | 0.42 ($p<10^{-4}$)*** | 0.25 ($p<10^{-4}$)*** | 0.54 ($p<10^{-4}$)*** | 0.35 ($p<10^{-4}$)*** |
| | Segment | Full | 0.67 ($p=0.02$)* | 0.67 ($p=0.02$)* | 0.59 ($p=0.04$)* | 0.54 ($p=0.06$) | 0.67 ($p=0.02$)* | 0.67 ($p=0.02$)* |
| | | Partial | 0.74 ($p=0.01$)* | 0.60 ($p=0.04$)* | 0.61 ($p=0.04$)* | 0.29 ($p=0.2$) | 0.74 ($p=0.01$)* | 0.60 ($p=0.04$)* |

\*: $p < 0.05$ ; \*\*: $p < 0.01$ ; and \*\*\*: $p < 0.001$ . The significant Spearman correlation coefficient ( $r_s$ ) are highlighted in gray. For the partial correlation, the neurite density index (NDI) and fractional anisotropy (FA) were included as covariates.

### Relevance to cortical myelin content

As illustrated in Fig. 5B, the length scaling coefficients were found to be negatively correlated with cortical myelin content (across voxels: HCP, *r_s_* = -0.59, *p* < 10^-4^; UKB, *r_s_* = -0.59, *p* < 10^-4^; across segments: HCP, *r_s_* = -0.76, *p* = 0.007; UKB, *r_s_* = -0.71, *p* = 0.01), suggesting a biological link between cortical myeloarchitecture and interhemispheric communication. After controlling for the NDI and FA, the partial Spearman correlations remained significant at both voxel and segment level (Table 1).

### Relevance to cortical expansion

To directly assess its evolutionary and developmental relevance, we investigated whether the length scaling of callosal fibers was related to the evolutionary and developmental expansion of their connected cortical regions, i.e., i) cortical expansion in humans relative to macaques and ii) cortical expansion in human adults relative to human infants. As described above, we assigned the mean cortical expansion index from the connected cortical region to each mCC voxel/segment. As shown in Fig. 5C-D, the mCC patterns of callosal fiber length scaling aligned well with those for the cortical expansion index during evolution (across voxels: HCP, *r_s_* = 0.75, *p* < 10^-4^; UKB, *r_s_* = 0.55, *p* < 10^-4^; across segments: HCP, *r_s_* = 0.75, *p* = 0.008; UKB, *r_s_* = 0.76, *p* = 0.006) and during development (across voxels: HCP, *r_s_* = 0.48, *p* < 10^-4^; UKB, *r_s_* = 0.42, *p* < 10^-4^; across segments: HCP, *r_s_* = 0.67, *p* = 0.02; UKB, *r_s_* = 0.67, *p* = 0.02). After controlling for the NDI and FA, the partial Spearman correlations remained significant at both voxel and segment level (Table 1).

### Callosal fibers of length over-scaling or under-scaling with brain size

Theoretically, a callosal fiber length coefficient of 1/3 indicates iso-scaling with brain size. In contrast, coefficients > 1/3 indicate that callosal fiber length scales more with greater brain size (i.e., over-scaling), and coefficients < 1/3 indicate that callosal fiber length scales less with brain size (i.e., under-scaling). As illustrated in Fig. 6A, significant mCC clusters (false discovery rate, FDR corrected *p* < 0.05) existed for both over-scaling and under-scaling. The overlapping regions between the clusters from the two datasets were designated as the final clusters; one large cluster exhibited over-scaling, and two small clusters demonstrated under-scaling (Fig. 6B). The over-scaling cluster covered most of the anterior half of the mCC, which mainly connects the prefrontal cortices between the two hemispheres. In contrast, the first under-scaling cluster was located on the splenium, mainly connecting the precuneus and superior parietal lobule; the other under-scaling cluster was on the posterior body, mainly connecting the paracentral lobule and precentral gyrus (Fig. 6C).

**Fig. 6.**
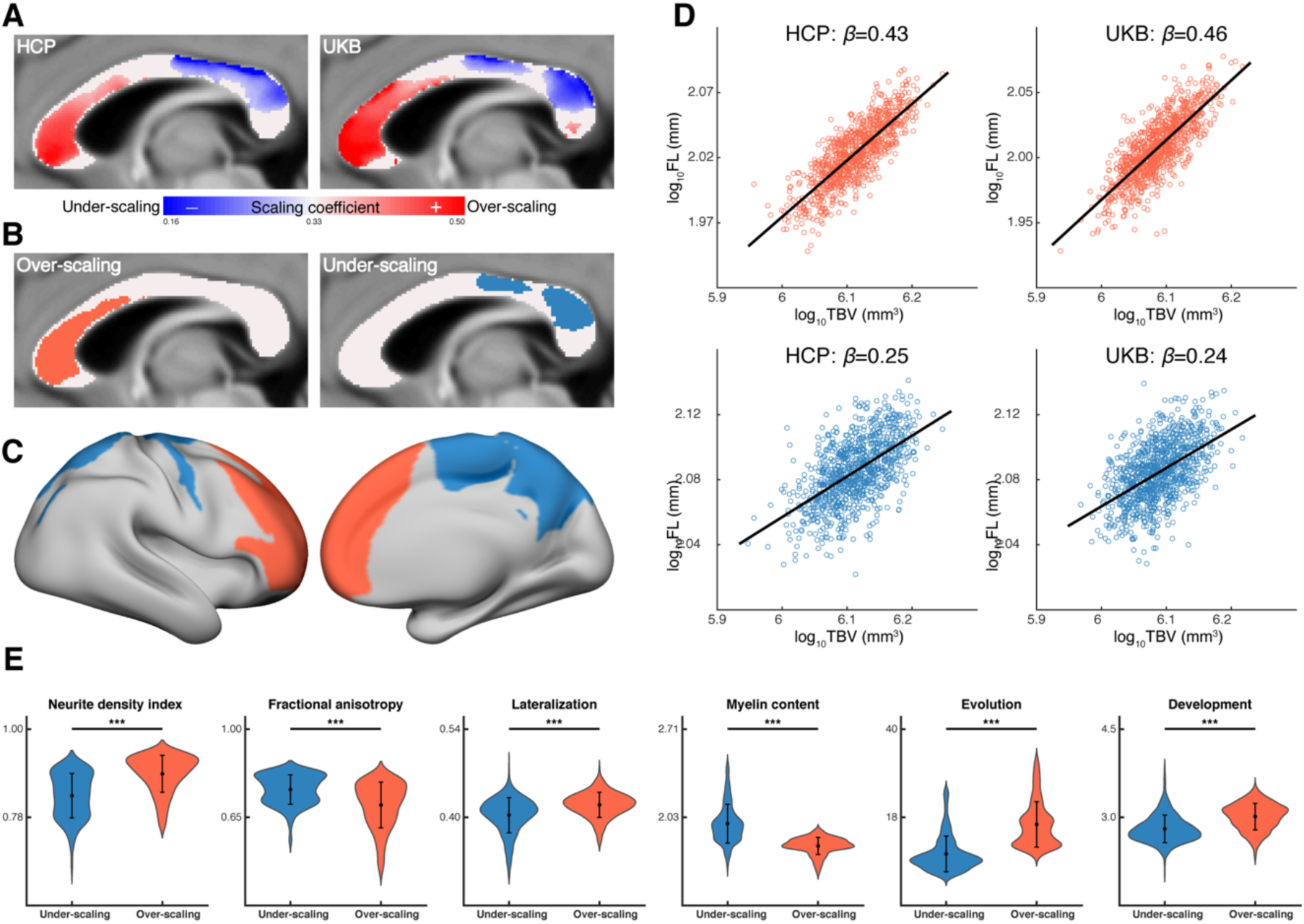
Callosal fibers of length over-scaling or under-scaling with brain size. (A) The midsagittal CC areas showing significant fiber length over-scaling or under-scaling. (B) The conjunction midsagittal CC area of length over-scaling or under-scaling between the HCP and UKB datasets. Left panel: over-scaling clusters. Right panel: under-scaling clusters. (C) The cortical regions connected by the callosal fibers passing through the over-scaling and under-scaling clusters. (D) Scatter plots of the logarithm of fiber length and the logarithm of TBV for the over-scaling (i.e., significant beta > 0.33) and under-scaling (i.e., significant beta < 0.33) clusters. (E) The differences between the over-scaling and under-scaling clusters in the two white matter microstructural measures and four cortical measures above. The mean (black dot) and standard deviation are overlaid on the violin map. ***: *p* < 0.001. Orange: over-scaling clusters. Blue: under-scaling clusters.

As shown in Fig. 6E, the callosal fibers of over-scaling showed greater NDI value (*t(826)* = 15.1, *p* < 10^-4^) and smaller FA value (*t(826)* = -10.9, *p* < 10^-4^), as well as stronger functional lateralization (*t(8691)* = 31.0, *p* < 10^-4^), less myelin content (*t(8691)* = -67.6, *p* < 10^-4^), and larger evolutionary and developmental expansion (*t(8691)* = 67.6, *p* < 10^-4^; *t(8691)* = 42.2, *p* < 10^-4^) in their connected cortical regions, compared with the callosal fibers of under-scaling. These results are compatible with the observed correlations of callosal fiber scaling with respect to the fiber composition and cortical measures described above.

Finally, our validation analyses confirmed the minimal influence of the procedure of mapping the cortical topography of mCC voxels or segments on our results presented above (Table 1 and Fig. 7).

**Fig. 7.**
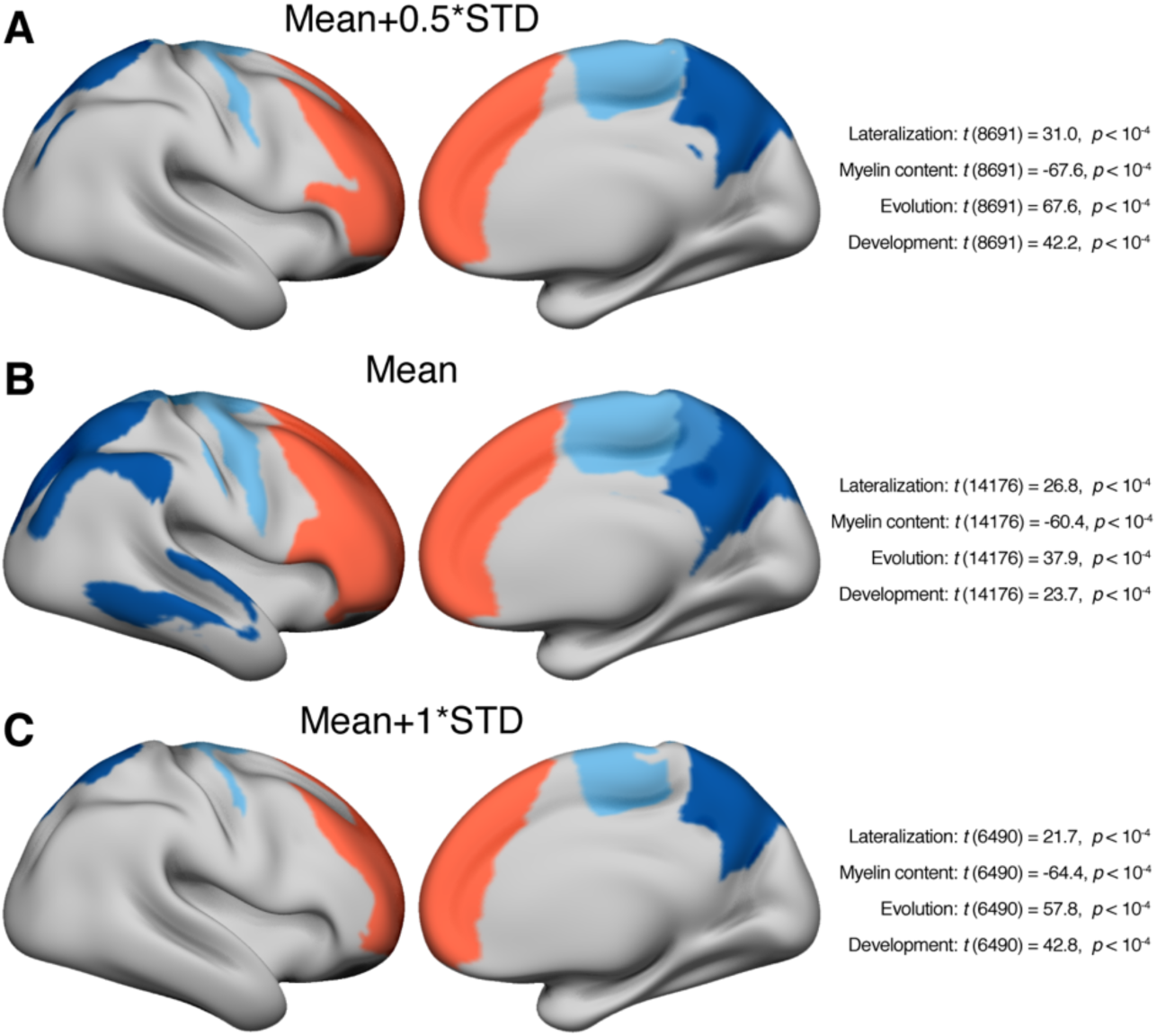
Cortical topography of the mCC clusters showing fiber length over-scaling and under-scaling. As shown in Fig. 6, there is one over-scaling mCC cluster (red) and there are two under-scaling mCC clusters (blue). To extract the binary cortical topography map, a threshold was applied to the group streamline count surface map. (A) The main threshold was defined as the mean + 0.5*STD of the group streamline count value across the entire surface (i.e., Fig. 6C). (B) The threshold was calculated as the mean of the group streamline count value across the entire surface. (C) The threshold was defined as the mean + 1*STD of the group streamline count value across the entire surface. Under each threshold, the differences between the over-scaling and under-scaling clusters of the functional lateralization index, of cortical myelin content, of evolutionary cortical expansion and of developmental cortical expansion are listed on the right side. The comparison showed consistent statistical results with our main text, indicating a limited impact of the cortical mapping threshold on the results.

## Discussion

Our present study revealed substantial variation in length scaling with brain size among callosal fibers, ranging from under-scaling to iso-scaling to over-scaling, replicated in two large healthy cohorts. Fiber length under-scaling was mainly observed in callosal fibers connecting the precentral gyrus and parietal cortices, whereas over-scaling mainly involved callosal fibers connecting the prefrontal cortices. Notably, the variation in such length scaling was biologically meaningful: the length scaling of callosal fibers significantly correlated with underlying fiber composition, as well as with human functional lateralization, myelin content, evolutionary and developmental expansion of the fiber-connected cortical regions. Validation analyses confirmed excellent reproducibility and robustness of these observations.

### Nonuniform length scaling of callosal fibers

The scaling patterns of GM features and midsagittal callosal size with brain size have been repeatedly studied, either across species or across human individuals (Jäncke et al., 1997; Rilling and Insel, 1999; Im et al., 2008; Herculano-Houzel, 2012; Mota and Herculano-Houzel, 2015; Reardon et al., 2018). In contrast, very few studies have been devoted to the scaling of WM fiber length with brain size, possibly due to technical difficulties in directly measuring fiber length *in vivo*. Our currently observed length scaling of callosal fibers and its biological relevance proved the overall feasibility and importance of studying WM fibers’ length scaling.

Notably, our current fiber length was estimated from dMRI tractography, so far the only technique of reconstructing WM fiber *in-vivo.* This widely-used technique however is inherently limited, showing errors and biases in resultant fiber streamlines particularly for long range WM tracts (Reveley et al., 2015; Donahue et al., 2016). To minimize these errors, the present study adopted high-angular resolution dMRI data, cutting-edge local orientation modeling, and tractography algorithm, i.e., the requirements for the most reliable fiber tracking. Moreover, we performed extensive validation analyses on our callosal tractography. While most validation results provided support for the overall validity of our callosal tractography, a few validation results suggested the existence of tractography biases. For example, the observed dispersion of callosal connected contralateral regions from the lateral temporal cortex (Fig. 3) likely reflects a systematic tendency or bias of callosal streamlines to avoid trajectories to the temporal lobe. In addition, the validity of our callosal tractography could be challenged to some extent by the lack of correlation between interhemispheric structural connectivity and functional connectivity (Fig. S6). Given these methodological limitations, the observed variation of length scaling among callosal fibers should be interpreted cautiously.

The substantial variation in length scaling suggests a change in the relative length among callosal fibers in larger brains. Such length reorganization in larger brains is unlikely confined to the CC but extrapolates to other WM fibers across the entire brain. On the other hand, previous studies have demonstrated fiber diameter/myelination reorganization in larger brains (Wang et al., 2008; Phillips et al., 2015). Thus, larger brains choose to jointly adjust the two critical determinants of conduction delay, i.e., fiber diameter/myelination and length, to reach an optimal fiber composition of the entire brain to achieve structural and functional efficacy best. Such a strategy is theoretically advantageous compared to solely adjusting either fiber diameter/myelination or length. The joint adjustment strategy offers a larger search space for the optimal fiber reorganization solution by providing two-parameter dimensions. Notably, the median or mean diameter of callosal fibers did not significantly differ between species or human individuals, although the largest diameter of callosal fibers did increase in species with larger brains (Olivares et al., 2001; Wang et al., 2008). Therefore, larger brains seem to adjust the relative length across the CC more than they adjust the fiber diameter/myelination. This particular adjustment model might be specific to the CC, and its underlying mechanisms deserve further investigation.

Callosal fiber density and diameters also exhibited significant variation in the mCC, as revealed by human histology (Aboitiz et al., 1992). Callosal fibers with larger diameters mainly connect sensorimotor cortices, and small diameters connect association cortices (Caminiti et al., 2013). This particular pattern reflects functional differences in cross-hemisphere communication among different types of cortical areas; sensorimotor functions require rapid interhemispheric information integration, and higher cognitive functions in association cortices work well with slow interhemispheric information transfer. In the present study, one of our observed under-scaling clusters was around the mCC posterior body, mainly connecting the sensorimotor cortex; the only cluster of over-scaling predominantly connected the prefrontal lobe—the primary association cortex. Therefore, the requirement of rapid communication between bilateral sensorimotor areas is accomplished by both larger fiber diameter and under-increased fiber length in larger brains. Smaller fiber diameter and an overly increased fiber length contribute to slower interhemispheric communication between bilateral association cortices. The rapid interhemispheric communication for low-level cortices but the slow interhemispheric communication for high-level cortices may reflect a general principle of the functional organization across species and individuals.

### Biological relevance of callosal fiber length scaling

The present study used two dMRI parameters to estimate the spatial pattern of mCC fiber composition *in-vivo*. Notably, in contrast to the FA, the NDI showed a strong correlation with the influential Aboitiz’s histological fiber density, so far the golden standard for such a measure (Aboitiz et al., 1992). This strongly verifies that the NDI can sensitively reflect axonal density in uncrossing areas (e.g., mCC), supporting relevant applications for this particular imaging parameter. Putatively, lower fiber density comes with a larger local fiber diameter on the mCC. The mCC FA did not correlate with histological fiber density and may reflect the myelination degree (Chang et al., 2017). The observed length scaling’s positive correlation with NDI and negative correlation with FA suggested that callosal fibers of less length scaling (i.e., smaller increase in fiber length) had lower density, larger diameter, and were myelinated. These empirical pieces of evidence support that smaller increase in fiber length, larger fiber diameter, and increased myelination are simultaneously implemented to jointly facilitate rapid interhemispheric communications of particular cortical regions in larger brains. On the other hand, the association between these callosal fiber characteristics may relate to cost control of fiber reorganization in larger brains: to save physical space and energy consumption, highly myelinated callosal fibers of large diameter tend to minimize their length increase as possible. Future studies are encouraged to relate the length scaling with other nontrivial callosal measures, e.g., midsagittal callosal thickness (Park et al., 2011).

Interhemispheric communication efficacy has long been considered a contributing factor to the emergence of functional lateralization. According to Ringo’s influential hypothesis, the excessive callosal conduction delay of larger brains leads to functional lateralization (Ringo et al., 1994). In contrast, functional lateralization may arise due to inter-hemispheric inhibition through callosal fibers (Cook, 1984). Measurements of the mCC area or dMRI parameters have been repeatedly used as a proxy for callosal connectivity to test these hypotheses (Bloom and Hynd, 2005; van der Knaap and van der Ham, 2011), with contrasting results. Both positive and negative correlations between callosal connectivity and functional lateralization have been reported (Josse et al., 2008; Karolis et al., 2019), supporting both hypotheses. By directly measuring callosal length, our observations suggested that more scaling of callosal fiber length (i.e., a relatively greater conduction delay in larger brains) is accompanied by stronger functional lateralization, therefore favoring Ringo’s hypothesis but not the inhibition hypothesis. Ringo’s original hypothesis did specify that an excessive interhemispheric conduction delay, which leads to functional lateralization, is mainly caused by increased callosal fiber length due to brain size expansion. Our observed positive correlation of callosal length scaling with functional lateralization provides more direct support for this influential hypothesis.

The association of the callosal fiber length scaling with cortical expansion during evolution and postnatal development provides direct support for evolutionary and developmental contributions to scaling variation. For a cortical region, less demand for rapid interhemispheric communication (as reflected by greater length scaling) and greater expansion of the cortical area are likely rooted in the same motivations for the entire brain’s optimal functional efficiency. As revealed previously, expansion of the cortical area across human individuals also showed under-scaling, iso-scaling, over-scaling with greater brain size (Reardon et al., 2018). The under- and over-expansion of cortical regions with greater brain size were attributed mainly to under- or over-increased cortical folding (Zilles et al., 2013). Notably, cortical expansion can be achieved by modifying the degree of cortical folding, surface outward degree, or both. Given the U shape of callosal fibers, their length scaling can partly represent their connected cortical regions’ outward surface degree. Therefore, the observation of callosal fiber length under- or over-scaling provides evidence for the contributing role of the surface outward degree in under- or over-expansion of the cortical area across individuals or during evolution and development. However, for specific regions, the extent that the changes in cortical folding degree or outward surface degree contribute to the evolutionary and developmental expansion of the cortical area remains elusive.

Nevertheless, biological mechanisms underlying the association of the length scaling with the nontrivial callosal and cortical measures remain unclear. It might relate to the shared genetic or environmental factors. At the neural level, causal relationships could exist among these fiber and cortical characteristics. Animal studies are warranted in the future in order to evaluate these speculations.

In conclusion, the current findings of callosal fiber variation in length scaling with brain size provide direct empirical evidence for length reorganization among WM fibers in larger brains. These observations also highlight the interactions of evolutionary and developmental constraints with interhemispheric communication.

## Acknowledgments

We thank Yanchao Bi for valuable discussions. This work was supported by the National Science Foundation of China [8217071142, 82021004, G.G.]. Michel Thiebaut de Schotten has received funding from the European Research Council (ERC) under the European Union’s Horizon 2020 research and innovation programme (grant agreement no. 818521). Data were provided by the Human Connectome Project, WU-Minn Consortium (Principal Investigators: David Van Essen and Kamil Ugurbil; 1U54MH091657) funded by the 16 NIH Institutes and Centers that support the NIH Blueprint for Neuroscience Research; and by the McDonnell Center for Systems Neuroscience at Washington University. UK Biobank brain imaging was funded by the UK Medical Research Council and the Wellcome Trust.

## Supplementary figures

**Supplementary Figure 1.**
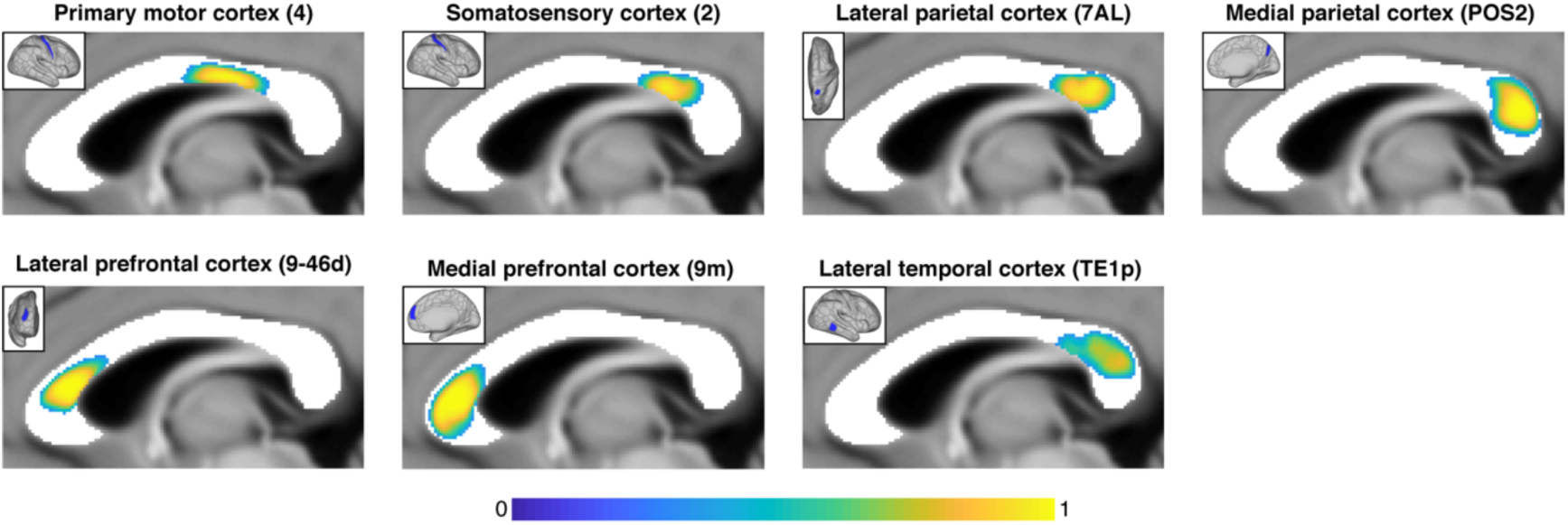
Callosal streamlines’ passing location on the mCC for selected cortical regions. The resultant group mCC pattern maps are displayed for selected HCPMMP regions on the right hemisphere (blue in solid box): area 4 on the primary motor cortex, area 2 on the somatosensory cortex, area TE1p on the lateral temporal cortex, area 7AL on the lateral parietal cortex, area POS2 on the medial parietal cortex, area 9-46d on the lateral prefrontal cortex, and area 9m on the medial prefrontal cortex. The HCPMMP regional abbreviation is included in the bracket.

**Supplementary Figure 2.**
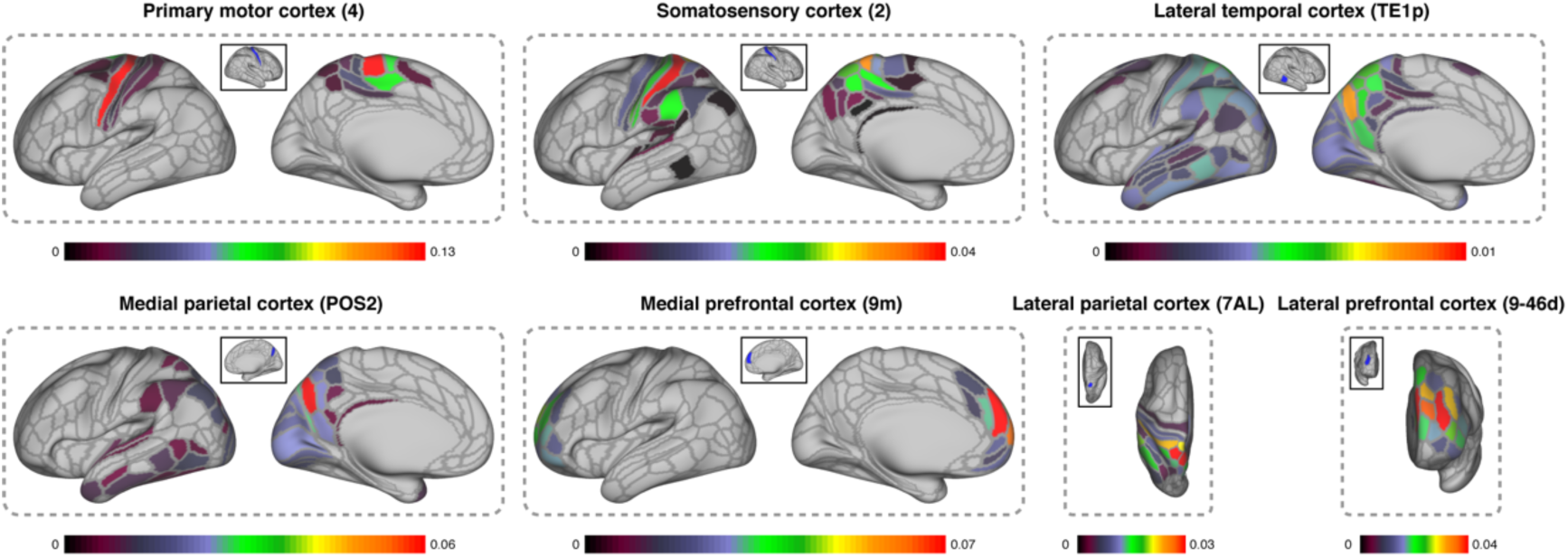
Major callosal connections to the contralateral hemisphere for selected cortical regions. For each cortical region, the top strongly connected regions on the contralateral hemisphere are displayed, which together account for at least 80% of total fraction of streamline (FSe) value from this region. The group FSe maps are displayed for selected HCPMMP regions on the right hemisphere (blue in the solid box): area 4 on the primary motor cortex, area 2 on the somatosensory cortex, area TE1p on the lateral temporal cortex, area 7AL on the lateral parietal cortex, area POS2 on the medial parietal cortex, area 9-46d on the lateral prefrontal cortex, and area 9m on the medial prefrontal cortex. The HCPMMP regional abbreviation is included in the bracket. The color bar indicates the FSe value from the selected cortical region.

**Supplementary Figure 3.**
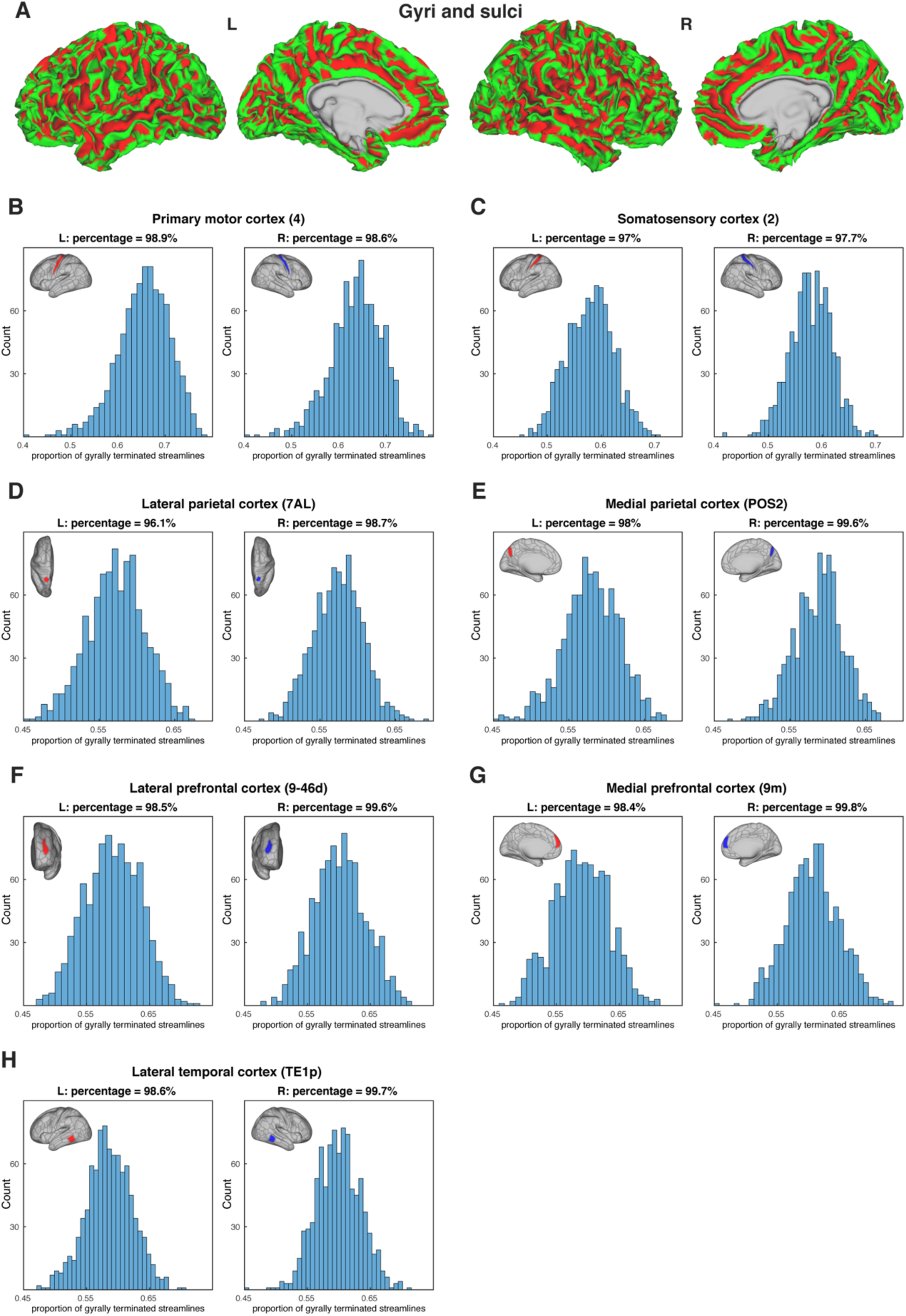
The proportion of gyrally terminated streamlines in the contralateral hemisphere for selected cortical regions. (A) The gyral (green) and sulcal (red) mask from one example subject. The histograms of proportion of gyrally terminated streamlines are displayed for selected HCPMMP regions: (B) area 4 on the primary motor cortex, (C) area 2 on the somatosensory cortex, (D) area 7AL on the lateral parietal cortex, (E) area POS2 on the medial parietal cortex, (F) area 9-46d on the lateral prefrontal cortex, (G) area 9m on the medial prefrontal cortex, and (H) area TE1p on the lateral temporal cortex. Red: cortical region on the left hemisphere; Blue: cortical region on the right hemisphere. The HCPMMP regional abbreviation is included in the bracket.

**Supplementary Figure 4.**
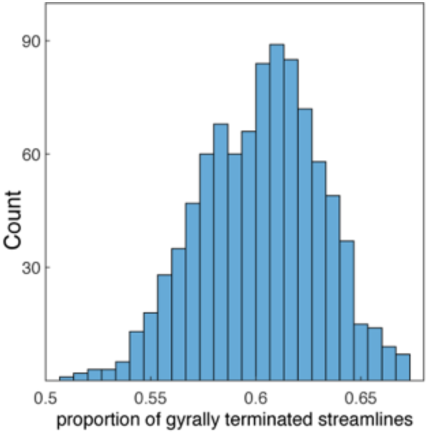
The proportion of gyrally terminated streamline for all callosal streamlines.

**Supplementary Figure 5.**
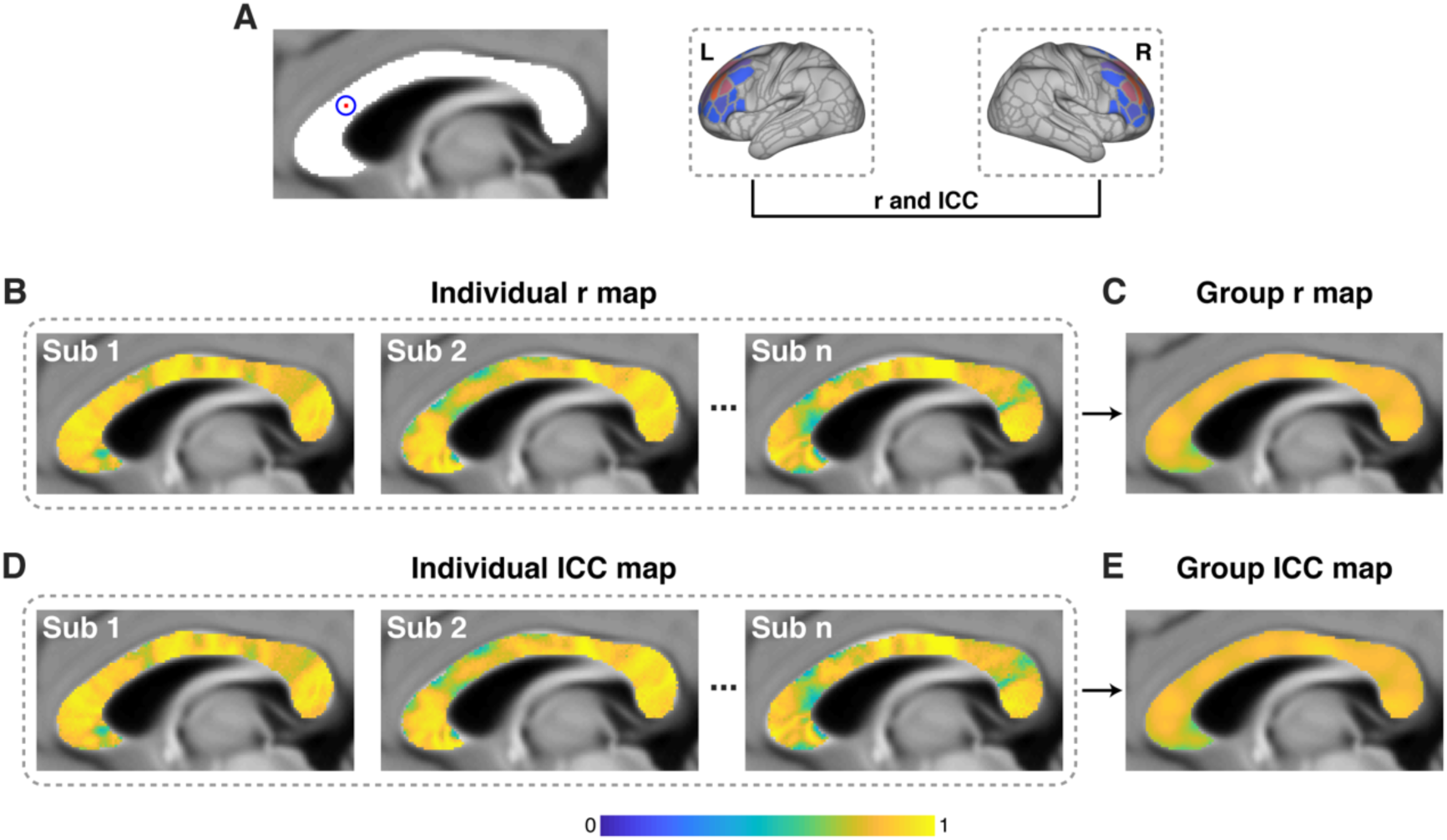
Spatial similarity of passing streamline termination pattern between the two hemispheres. (A) Schematic diagram of calculating *r* and *ICC* of termination distribution between left and right hemispheres for one mCC voxel. Individual *r* map (B) and *ICC* map (D) of the mCC. Group *r* map (C) and *ICC* map (E) of the mCC.

**Supplementary Figure 6.**
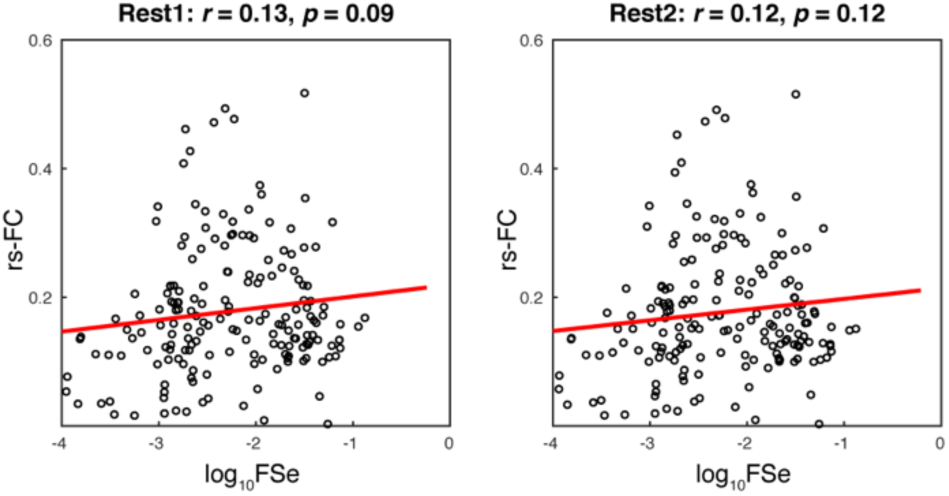
Correlation between structural connectivity and functional connectivity. Scatter plots of structural connectivity (FSe) and resting-state functional connectivity (rs-FC, Pearson correlation) across the 180 HCPMMP homotopic regional pairs. Left: Rest1; Right: Rest2.

